# To CRUNCH or not to CRUNCH: Task Difficulty Affects Functional Brain Reorganisation during Visuospatial Working Memory Performance in Premanifest Huntington’s Disease

**DOI:** 10.1101/459180

**Authors:** Maria V. Soloveva, Sharna D. Jamadar, Dennis Velakoulis, Govinda Poudel, Nellie Georgiou Karistianis

## Abstract

Presymptomatic Huntington’s disease (pre-HD) individuals tend to increase functional brain activity to compensate for HD-related brain anomalies. We used a quantitative model of compensation, known as the CRUNCH (Compensation-Related Utilization of Neural Circuits Hypothesis) to explicitly characterise compensation in pre-HD. We acquired functional magnetic resonance imaging (fMRI) data (*n* = 15 pre-HD; *n* = 15 controls) during performance of an 18-minute fMRI visuospatial working memory task with low, intermediate-1, intermediate-2, and high memory loads. Consistent with the CRUNCH prediction, pre-HD individuals showed decreased fMRI activity in left intraparietal sulcus at high memory load, compared to healthy controls who showed increased fMRI activity in left intraparietal sulcus at high memory load. Contrary to the other CRUNCH prediction, the pre-HD group did not show compensatory increase in fMRI activity at lower levels of memory loads in left intraparietal sulcus. Our findings provide partial support for the validity of CRUNCH in pre-HD.

**Highlights:**

- Visuospatial working memory deficits in pre-HD occur 25 years prior to predicted disease onset
- Task demands differentially affect fMRI activity in left intraparietal sulcus
- CRUNCH can partially apply in Huntington’s disease

## 1. Introduction

Huntington’s disease (HD) is a fatal genetic neurodegenerative disorder caused by a cytosine-adenine-guanine (CAG) repeat expansion in the Huntingtin gene (*HTT*) on chromosome 4 (Macdonald et al., 1993). During the premanifest stage of HD (pre-HD), significant brain changes have been documented as early as 20 years prior to clinical diagnosis. These include cortical atrophy (Beglinger et al., 2005;Domínguez et al., 2013;Henley et al., 2009), sub-cortical atrophy (Dominguez et al., 2013;Dominguez et al., 2016; Georgiou-Karistianis, Scahill, Tabrizi, Squitieri, & Aylward, 2013a; Georgiou-Karistianis et al., 2013b; Tabrizi et al.,2013), compromised white matter integrity (Domínguez et al., 2013; Poudel et al., 2015a; Poudel et al., 2014a), impaired resting-state functional connectivity (Harrington et al., 2015; Poudel et al., 2014b), as well as functional magnetic resonance imaging (fMRI) task-related activity changes (Dominguez et al., 2017; Georgiou-Karistianis et al., 2013c; Georgiou-Karistianis et al., 2014; Gray et al., 2013; Kloppel et al., 2015; Kloppel et al., 2009; Malejko et al., 2014; Poudel et al., 2015b; Wolf, Vasic, Schonfeldt-Lecuona, Landwehrmeyer, & Ecker, 2007). Despite these widespread brain anomalies, pre-HD individuals are able to maintain a normal level of cognitive functioning required to undertake routine day-to-day activities.

Pre-HD individuals typically exhibit increased fMRI activity and recruit additional brain regions during task performance to maintain a comparable level of behavioural performance to healthy controls. Increased and/or additional fMRI activity is evident when performing tasks, such as verbal working memory (Kloppel et al., 2009; Kloppel et al., 2015; Wolf et al., 2007), spatial working memory (Georgiou-Karistianis et al., 2013c; Poudel et al., 2015b), set-shifting (Gray et al., 2013), time discrimination (Paulsen et al., 2004), sequential finger tapping (Scheller et al., 2013), and reward processing (Malejko et al., 2014). The pattern of fMRI overactivation coupled with preserved cognitive and motor performance has been often interpreted as compensation. However, this ‘informal’ definition is problematic. For example, fMRI over-activation can be also attributed to: (1) medication effects (Ianetti & Wise, 2007); (2) dedifferentiation (Baltes & Lindenberger, 1997); (3) an inability of the brain to inhibit fMRI activity not related to the task being performed (Cox et al., 2015); and/or (4) individual differences in the physiology of the fMRI blood oxygenation level dependent (BOLD) response (Iannetti & Wise, 2007). Most importantly, HD is a progressive disease with widespread brain pathology, e.g., cortical and sub-cortical atrophy, changes in neurovascular coupling (Hua, Unschuld, Margolis, Zijl, & Ross,2014), and it is therefore difficult to disentangle whether fMRI over-activation reflects true compensation in pre-HD or whether it is a product of HD-related pathological processes.

The HD community has started to advocate the need to better characterise increased and/or additional fMRI activity in HD based on an *explicit* model/definition of compensation (Gregory et al., 2018; Gregory et al., 2017; Kloppel et al., 2015; Soloveva, Jamadar, Poudel, & Georgiou-Karistianis, 2018). For example, Gregory et al. (2017) proposed a longitudinal model of compensation to explicitly characterise fMRI over-activation in neurodegenerative disorders. According to the model, compensatory response in HD is defined as ‘concave-down’ longitudinal changes of fMRI activity and behavioural performance in the presence of linear decreases of brain volume over time. Consistent with the model, Gregory et al. (2018) demonstrated a pattern of longitudinal compensation in the cognitive (dorsolateral prefrontal cortex) and motor networks (premotor cortex) in pre-HD individuals and early HD patients, compared with healthy controls. Although these operational definitions of compensation could be an important step forward the characterisation of compensation in pre-HD, they fail to acknowledge that pre-HD individuals can compensate even in the absence of improved cognitive performance (Zarahn, Rakitin, Abela, Flynn, & Stern, 2007). Strictly speaking, if a compensatory mechanism was not in place, pre-HD individuals would either perform as effectively as heathy controls or poorer. As such, other models of compensation to characterise compensation in pre-HD should be considered.

Other such models including the Hemispheric Asymmetry Reduction in OLDer Adults (HAROLD; Cabeza, 2002), Posterior-Anterior Shift in Ageing (PASA; Davis, Dennis, Daselaar, Mathias, & Cabeza, 2007), Scaffolding Theory of Ageing and Cognition (STAC; (Park & Reuter-Lorenz, 2009), Scaffolding Theory of Ageing and Cognition Revised (STAC-r; Reuter-Lorenz & Park, 2014), Model of Aged-Related Compensation (Cabeza & Dennis, 2013), and Compensation Related Utilization of Neural Circuits Hypothesis (CRUNCH; Reuter-Lorenz & Cappell, 2008) have been proposed to clarify whether increased fMRI activity reflects a beneficial compensatory response, rather than age-related brain anomalies. In a recent review (Soloveva et al., 2018), we critically reviewed each of the models in the context of HD functional neuroimaging literature, and suggested the CRUNCH (Reuter-Lorenz & Cappell, 2008) may be the most relevant model of compensation to test in HD (see Figure 1 (A)). Importantly, CRUNCH offers a quantifiable and explicit test for the link between fMRI activity and behavioural performance (Soloveva et al., 2018). In particular, CRUNCH proposes that fMRI activity increases as task difficulty increases (usually at lower task demands) to compensate for task difficulty. However, when individuals reach a critical point (the ‘CRUNCH’ point), where task difficulty exceeds their capacity (usually at higher task demands), fMRI activity and behavioural performance declines. CRUNCH postulates that older adults typically show increased fMRI activity at lower task demands (compensation overactivation effect). The model hypothesises that older adults reach the ‘CRUNCH’ point at intermediate levels of task demands because the compensatory mechanism is no longer effective, and results in decreased fMRI activation in older vs younger adults.

**Figure 1.**
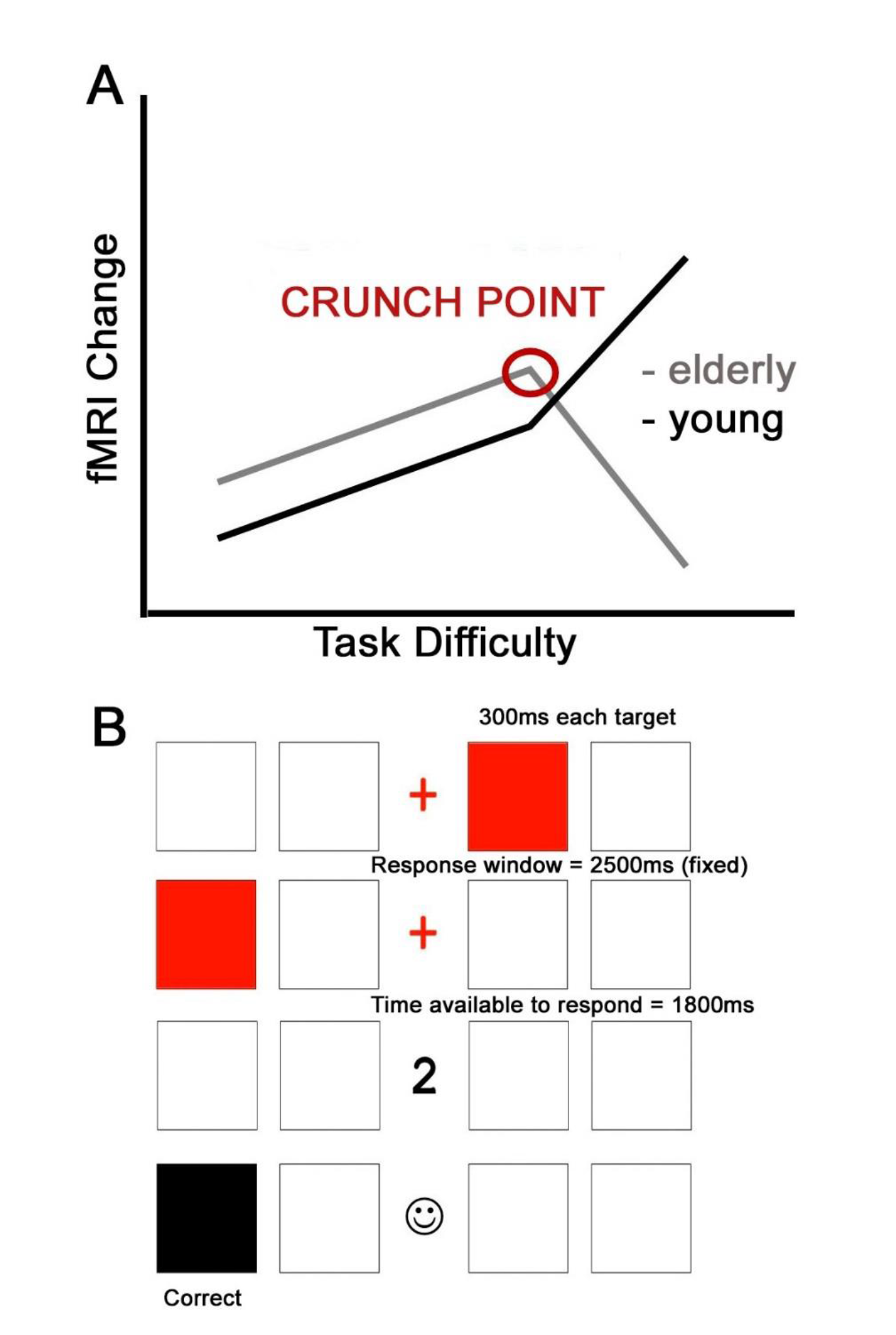
(A) Visual representation of the hypothesised relationship between fMRI activity and task difficulty as proposed by the CRUNCH model (Reuter-Lorenz & Cappell, 2008). As task difficulty increases, older adults show increased fMRI activity. However, once their resources are exceeded, they reach the ‘CRUNCH’ point, and their fMRI activity and behavioural performance declines. By comparison, young adults show greater fMRI activity at high task demands only. (B) Visual representation of the fMRI visuospatial working memory task. In this example, two boxes are highlighted (low level of difficulty). A cue (number ‘2’) prompts the participant to indicate which box was highlighted second in the sequence. Participants can respond as soon as they see the cue.

We used the CRUNCH model (Reuter-Lorenz & Cappell, 2008) to explicitly characterise compensation in pre-HD during visuospatial working memory, using a task that incremented in levels of difficulty (low, intermediate-1, intermediate-2, and high). This task was selected because visuospatial working memory is one of the earliest signs of cognitive deficits in pre-HD (Dumas et al., 2012) and no HD study to date has tested explicitly compensatory response in this cognitive domain. In accordance with the CRUNCH model, we hypothesised that: (a) pre-HD individuals would show increased fMRI activity at low memory loads; and (b) decreased fMRI activity at higher memory loads, compared to healthy controls.

## 2. Methods

### 2.1. Participants

The study was approved by the Monash University and Melbourne Health Human Research Ethics Committees and written informed consent was obtained from each participant in accordance with the Helsinki Declaration. A total of 33 participants were recruited for the study, consisting of 17 pre-HD individuals (*n* = 16 right-handed, *n =* 1 left-handed) and 16 (*n* = 15 right-handed; *n =* 1 left-handed) healthy controls. Three participants were excluded, including two pre-HD individuals and one healthy control on the basis of the following criteria: (1) failure to perform with ≥ 70% behavioural accuracy during the low task difficulty condition (*n* = 1 healthy control); (2) movement of greater than 3mm during the MRI scan (*n* = 1 pre-HD); and (3) claustrophobia (*n* = 1 pre-HD). The final sample therefore consisted of 30 participants, consisting of 15 pre-HD individuals (*n* = 14 right-handed; *n =* 1 left-handed) and 15 age- and gender-matched (*n* = 14 right-handed; *n =* 1 left-handed) healthy controls. Demographic, clinical and neurocognitive details for the 30 participants are included in Table 1.

**Table 1.** Demographic information, clinical measures and neurocognitive data.

| | Means ± SD | |
| --- | --- | --- |
|  | Controls | Pre-HD |
| n (sample sizes) | 15 | 15 |
| Gender (M:F) | 5:10 | 5:10 |
| Age | 35.60 ± 10.69 | 37.33 ± 10.82 |
| IQ estimate | 109.33 ± 6.85 | 109.07 ± 6.61 |
| UHDRS | N/A | 0.005 ± 0.02 |
| CAG repeats | N/A | 40.87 ± 2.23 |
| Disease burden score | N/A | 176.33 ± 75.88 |
| SDMT | 64.47 ± 11.07 | 55.20 ± 12.60 |
| BDI-II | 3.73 ± 4.35 | 3.52 ± 3.04 |
| MoCA | 28.13 ± 1.46 | 26.96 ± 1.47 |

SD, standard deviation; IQ, estimate of full scale IQ, National Adult Reading Test (NART-2) error score; UHDRS, motor subscale score, Unified Huntington's Disease Rating Scale (pre-HD, UHDRS ≤ 5); CAG, cytosine-adenine-guanine (number of repeats > 40 is full penetrance; Disease Burden Score (CAG-35.5)\*age; SDMT, Symbol Digit Modalities Test; BDI-II, Beck Depression Inventory score Version II; MoCA, Montreal Cognitive Assessment score.

No significant differences in IQ (NART-2; Nelson, Willison, & Owen, 1992) and BDI- II (Beck, Steer,& Brown, 1996) scores between pre-HD individuals and healthy controls were revealed by an independent-sample t-test. Age was not used as a covariate in the analysis as pre-HD individuals and healthy controls were age- and gender-matched a priori. Behavioural data were analysed using SPSS Statistics v22.

All individuals were free from: (1) severe psychiatric and neurological illness (other than a diagnosis of HD for the pre-HD group); (2) brain injury or stroke; (3) substance abuse disorder; (4) intellectual disability. Based on clinical examination all pre-HD individuals had UHDRS total motor score of ≤ 5 (Tabrizi et al., 2009). According to the formula by Langbehn, Brinkman, Falush, Paulsen, & Hayden, (2004) based on age and the number of CAG repeats, the mean pre-HD group’s estimated years to disease onset was 25.77 ± 12.79 years with a mean disease burden score of 176.33 ± 75.88 (Orth & Schwenke, 2011). Pre-HD individuals reported the following medications: serotonin-norepinephrine reuptake inhibitor (SNRI) (*n* = 1), selective serotonin and norepinephrine reuptake inhibitor (SSNRI) (*n* = 1), selective serotonin reuptake inhibitor (SSRI) (*n* = 2), and mirtazapine antidepressants (*n* = 1). In addition, one pre-HD individual was taking Clobazam for restless leg syndrome and one was taking thyroxine to maintain normal thyroid hormone levels, as well as an atypical antipsychotic to stabilise mood and cognition. Healthy controls reported taking preventive treatment for HIV (*n* = 1) and an ion supplement (*n* = 1).

All participants underwent neurocognitive and psychological tasks sensitive across disease stages (Tabrizi et al., 2012): the Montreal Cognitive Assessment (MoCA; (Nasreddine et al., 2005), the NART-2 (Nelson et al., 1992), the Symbol Digit Modalities Test (SDMT; Smith, 1991), and the Beck-Depression Inventory- II (BDI- II; Beck et al., 1996). Following neurocognitive and psychological testing, participants performed a ∼ 10-minute block of training on the visuospatial working memory task with low, intermediate-1, intermediate-2 and high difficulty conditions (Figure 1(B)). Each participant completed 10 trials per condition and was required to reach the accuracy cut-off of ≥ 70% at low level of difficulty. This was to ensure that all participants were able to complete the task in the MRI scanner and to minimise the effects of learning (e.g., stimulus-response mapping) on in-scanner task performance. Lastly, participants underwent a subsequent 1-hour structural and functional MRI (fMRI) scan.

### 2.2. fMRI visuospatial working memory experimental paradigm

The fMRI experimental paradigm is an 18-minute novel task to measure visuospatial working memory under varied memory loads (targets) between three blocks. The memory load parametrically varied from low (2 targets), intermediate-1 (3 targets), intermediate-2 (4 targets) and high memory load (5 targets). Low, intermediate-1, as well as intermediate-2 difficulty condition consisted of 31 trials each, and there were 37 trials per high difficulty condition. In this task, participants were required to store the sequence of red highlighted boxes in visuospatial working memory (2, 3, 4, or 5 targets) and after a delay (1000ms) to replicate it with button-presses as fast as possible. Each target was displayed for 300ms with a response to be made within 1800ms. Participants responded with four fingers (index and middle finger of each hand) which mapped spatially to the four presented boxes to indicate which box was highlighted 1st, 2nd, 3rd, 4th, or 5th in a sequence. The order of load conditions was not counterbalanced to avoid an asymmetric switch cost when switching between tasks of different difficulty (Monsell, Yeung, & Azuma, 2000). Primary outcome measures were mean reaction times (ms) and the mean number of errors per task difficulty condition.

### 2.3. Functional data acquisition

Structural and functional MRI images were acquired using a Siemens 3-Tesla Skyra MRI scanner with a 20-channel head coil located at Monash Biomedical Imaging, Melbourne, Australia. High-resolution T1- weighted images were acquired to characterise structural integrity with sub-millimeter resolution (192 slices, 1mm slice thickness, 1mm x 1mm x 1mm voxel size, TE = 2.07ms, TR = 2300ms, FoV = 256mm; flip angle 9°). Functional images were acquired during the task performance using T2*-weighted BOLD scans with a gradient echo-planar sequence using interleaved slice orientation (44 slices; 3mm slice thickness, 2.5mm x 2.5mm x 3mm voxel size, TE = 30ms; TR = 2550ms; FoV = 192mm, flip angle = 80°). Time of acquisition was 1 hour per individual.

### 2.4. Data Analysis

#### 2.4.1. Behavioural data analysis

Mean reaction times (RTs) and the mean number of errors were recorded for each participant for each level of difficulty. Group means and standard errors for both groups are presented in Figure 2. RT and number of errors were analysed using separate 2 Group (pre-HD, healthy controls) by 4 Level of Difficulty (low, intermediate-1, intermediate-2 and high) mixed model ANOVA.

**Figure 2.**
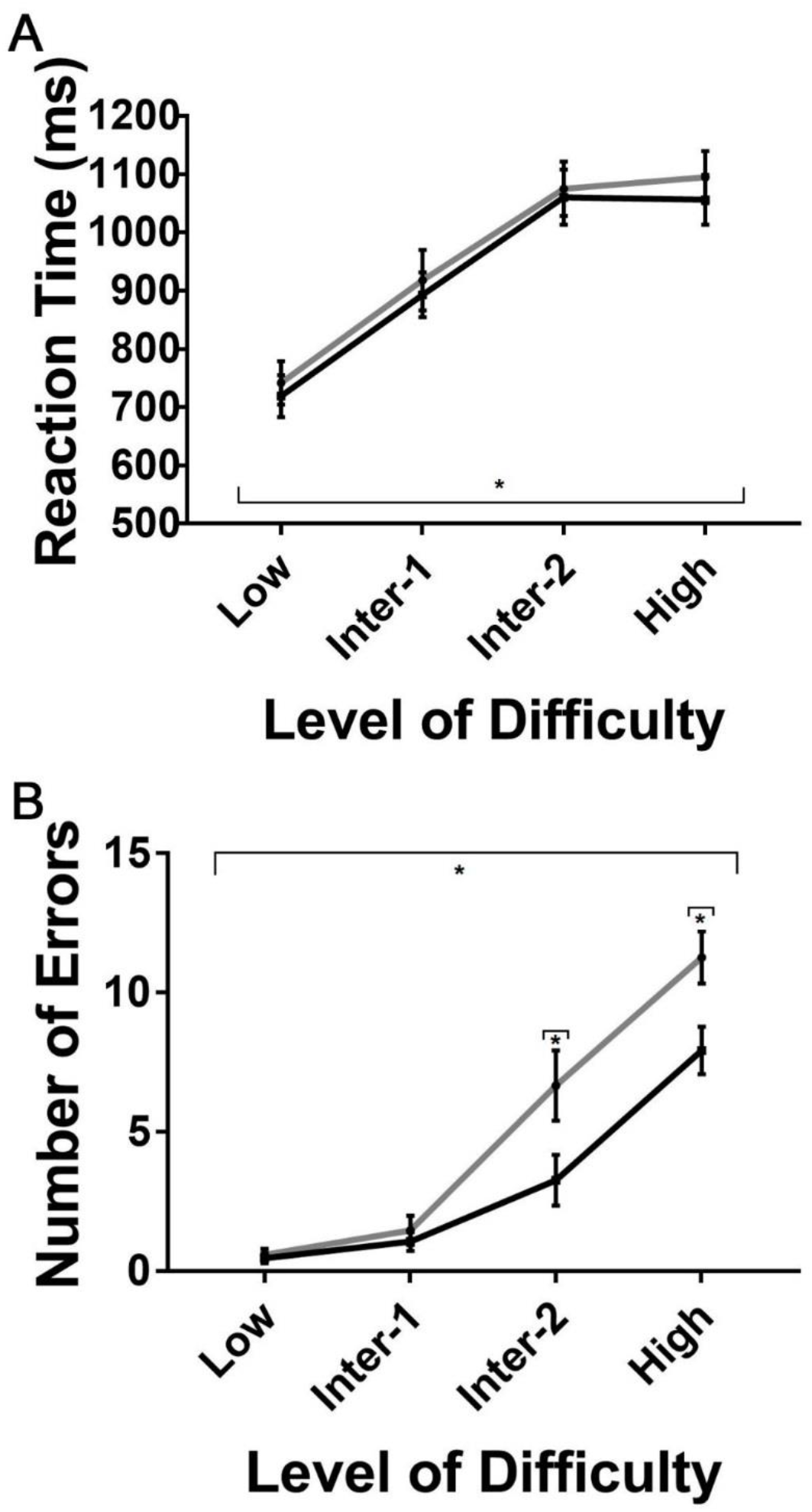
(A) Mean reaction times (ms) for pre-HD individuals and healthy controls for each difficulty level. The grey line corresponds to reaction times of pre-HD individuals. The black line corresponds to reaction times of healthy controls. Standard errors are represented in the figure by the error bars for each group for each difficulty level. Black asterisk denotes main effect of Level of Difficulty at *p* <.05. (B) Mean number of errors for pre-HD individuals and healthy controls for each difficulty level. The grey line corresponds to number of errors of pre-HD individuals. The black line corresponds to number of errors of healthy controls. Standard errors are represented in the figure by error bars for each group for each difficulty level. Black asterisks denote main effect of Group and the Group by Level of Difficulty interaction at *p* > .05.

#### 2.4.2. fMRI spatial pre-processing

Functional and structural MRI data were preprocessed and analysed using Statistical Parametric Mapping software (Wellcome Institute of Neurology at University College, London, UK; SPM12; http://www.fil.ion.ucl.ac.uk/spm/software/) implemented in MATLAB 8.0 (MathWorks, Natick, MA).

The first five scans of every fMRI run were discarded to account for the T1 saturation effect. The fMRI data underwent slice-timing correction followed by rigid-body realignment of all images to the middle slice to correct for head motion. Motion parameters were screened for each participant, and all participants showed less ≤ 3mm head motion, and was thus satisfactory. Subsequently, T1-weighted structural images were segmented into separate cerebrospinal fluid, white matter, and grey matter tissue probability maps. To improve robustness of the registration process (T1 images contain considerable amounts of non-brain tissue) (Fischmeister et al., 2013), we also edited scalp to remove non-brain areas before normalisation. The realigned fMRI scans were then co-registered to each individual’s structural T1-weighted skull-stripped image. The T1 image was then spatially normalised to Montreal Neurological Institute (MNI) space using non-linear transformation routines implemented in SPM12. The resulting transformation parameters were applied to the functional images. All images were inspected visually at each step to ensure adequate registration and normalisation. Finally,functional images were smoothed with a 6mm full width at high maximum isotropic Gaussian kernel and high pass filtered (128s).

#### 2.4.3. Volumetric Data Analysis

To ensure that there were no significant differences in brain volume between pre-HD and healthy controls, voxel-based morphometry analyses (VBM) were performed with SPM12 (Wellcome Institute of Neurology at University College, London, UK). Each participant’s unnormalised T1 structural image was segmented into grey matter (GM), white matter (WM) and cerebrospinal fluid (CSF) according to VBM protocol implemented as part of cat12 (http://www.neuro.uni-jena.de/cat/). The images were resliced with 1.0 by 1.0 by 1.0 mm^3^ voxels producing grey and white matter images. The resulting modulated and normalised GM and WM images were further smoothed with a 6mm full width at high maximum isotropic Gaussian kernel. Modulated and normalised VBM data were used for the group comparisons of GM and WM (i.e., comparisons of an absolute amount of tissue type within a region) (Ashburner & Friston, 2001). To identify whether there were significant differences between pre-HD individuals and healthy controls on measures of GM, WM and TIV, we performed an independent-sample *t*-test. No significant differences were revealed between pre-HD individuals and healthy controls on measures of GM, t(28) = -.08, *p =* .94, WM, t(28) = -.66, *p =* .52, and TIV, t(28) = -.62, *p =* .54. As such, we did not control for group differences in brain volume in the fMRI analyses.

#### 2.4.4. fMRI data analysis

Functional data were analysed within the framework of the General Linear Model (GLM) (Friston et al., 1995). At the first level, task-related fMRI signal changes were analysed for each level of difficulty (low, intermediate-1, intermediate-2, and high); incorrectly performed trials were entered as regressors of no interest and six motion parameters were included as nuisance regressors to account for residual effects of head motion. Contrast images for *low-correct-trials > baseline, intermediate-1 -correct-trials* > *baseline, intermediate-2- correct-trials* > *baseline,* and *high-correct-trials* > *baseline* were computed for each participant. Here, ‘baseline’ refers to implicit baseline; i.e., unmodelled variance in the GLM.

At the second level, contrast images for each contrast of interest were submitted to second-level one-sample t-test to analyse fMRI activation at whole-brain level, threshold of *p* <.001 (uncorrected), minimum cluster size of 50 voxels (*p* FWE corrected < .05 at cluster-level). To examine the effects of difficulty and group, we performed a region-of-interest (ROI) analysis. Parameter estimates (contrast values) were calculated for each ROI for *low-correct-trials > baseline, intermediate-1-correct-trials* > *baseline, intermediate-2-correct-trials* > *baseline*, and *high-correct-trials* > *baseline* contrasts. The ROI analysis was focused on the regions derived from a meta-analysis by Rottschy et al. (2012) which identified key brain regions involved in working memory function. These ROIs included left and right: (1) anterior insula; (2) inferior frontal gyrus; (3) posterior medial frontal cortex; (4) intraparietal sulcus; and (5) dorsolateral prefrontal cortex. In order to target the striatum, the main site of neuropathology in HD (Albin et al., 1992; Rajkowska, Selemon, & Goldman-Rakic, 1998), we also included left and right caudate and putamen (Georgiou-Karistianis et al., 2014). ROIs were created as a 10mm sphere centred on the MNI coordinates reported by Rottschy et al. (2012) and Georgiou-Karistianis et al. 2014 using MarsBar (Brett, Anton, Valabregue, & Poline, 2002). Parameter estimates for each ROI were assessed using separate 2 Group (pre-HD, healthy controls) by 4 Level of Difficulty (low, intermediate-1, intermediate-2, and high) mixed model ANOVA (SPSS v.22).

## 3. Results

### 3.1. Behavioural results

The mean RTs and the mean number of errors for each group and difficulty level are presented in Figure 2.

Results from the mixed-design ANOVA confirmed that there was a significant main effect of Level of Difficulty on RTs, F(2.33; 65.09) = 121.20, *p* <.001, indicating that both groups were slower with increased task difficulty (Figure 2(A)). The Group by Level of Difficulty interaction was not significant, F(2.33; 65.09) = .13, *p =* .91, indicating that the effect of task difficulty did not differ between groups. Lastly, there was no significant Group difference for RT, F(1, 28) = .21, *p =* .65.

Results from the mixed-design ANOVA for the number of errors revealed a significant main effect of Group, F(1, 28) = 5.34, *p =* .03, indicating that pre-HD individuals made more errors overall, compared with healthy controls (Figure 2(B)). The number of errors in pre-HD individuals and healthy controls significantly increased as a function of Level of Difficulty, F(1.94; 53.60) = 102.79, *p* < .001. Lastly, the Group by Level of Difficulty interaction was statistically significant, F(1.94; 53.60) = 4.79, *p* = .01. Post-hoc analyses revealed that pre-HD individuals made significantly more errors at intermediate-2, F(1, 28) = 4.78, *p* = .04 and high level of difficulty, F(1, 28) = 5.34, *p =* .03, compared to healthy controls. However, the number of errors was similar for pre-HD individuals and healthy controls at low, F(1, 28) = .24, *p* = .63 and intermediate-1 task level of difficulty, F(1, 28) = .42, *p =* .52.

### 3.2. fMRI Results

#### 3.2.1. Whole Brain Analysis: One-sample T-test

The whole-brain analysis (*all-correct-trials > baseline*) (thresholded with *p* < .001 at voxel level and *p* <.05 at FWE cluster-level corrected) averaged across groups and all four levels of difficulty is reported in Table 2 and Figure 3. The task activated a distributed network, including bilateral temporo-occipital, premotor-parietal, and medial fronto-parietal regions, as well as left and right putamen and right caudate.

**Table 2.**
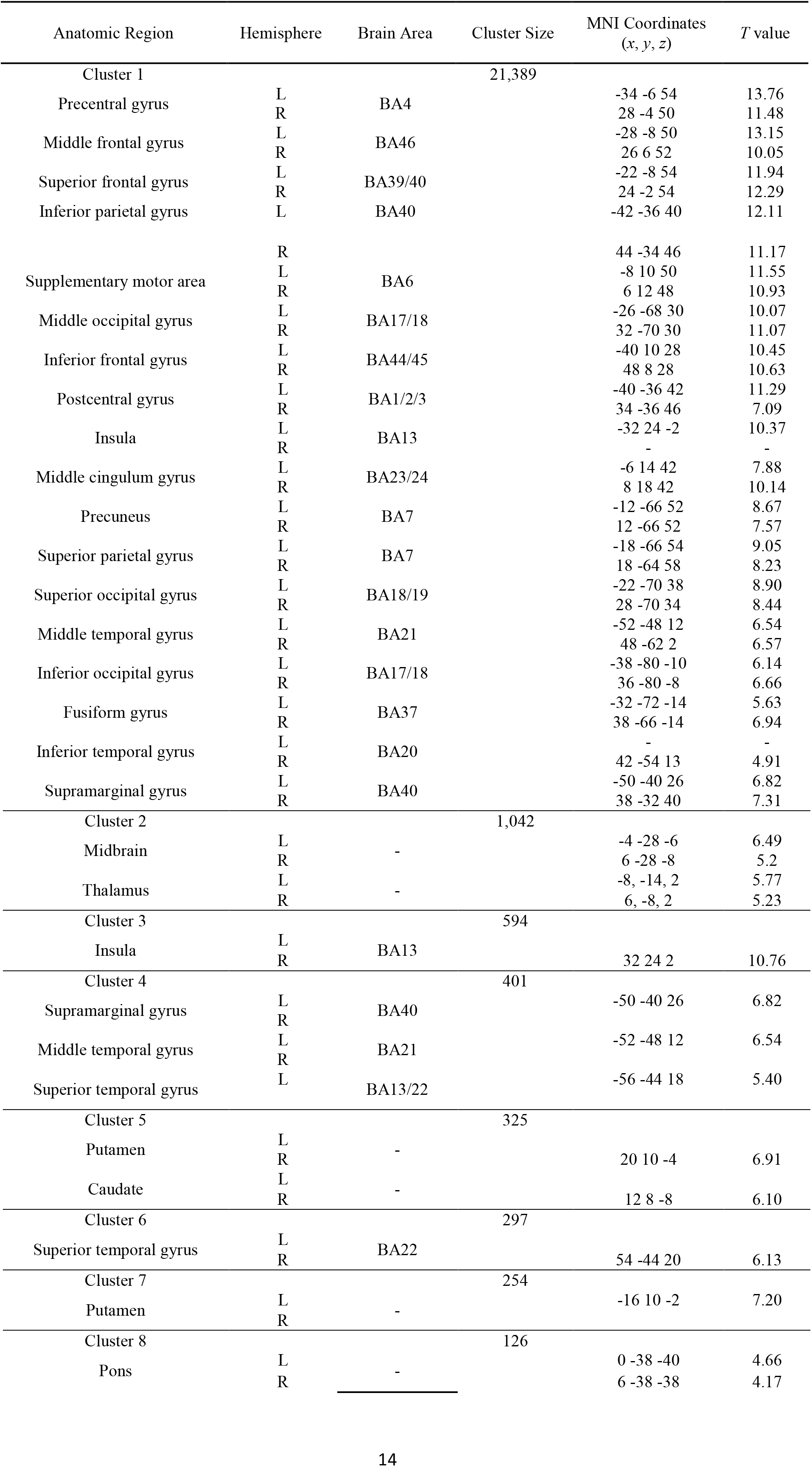
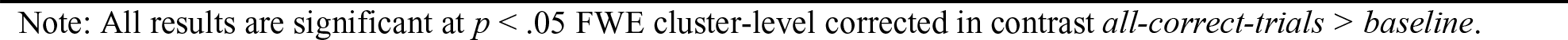
Brain regions showing significant fMRI activity in pre-HD and healthy controls during novel visuospatial working memory task.

**Figure 3.**
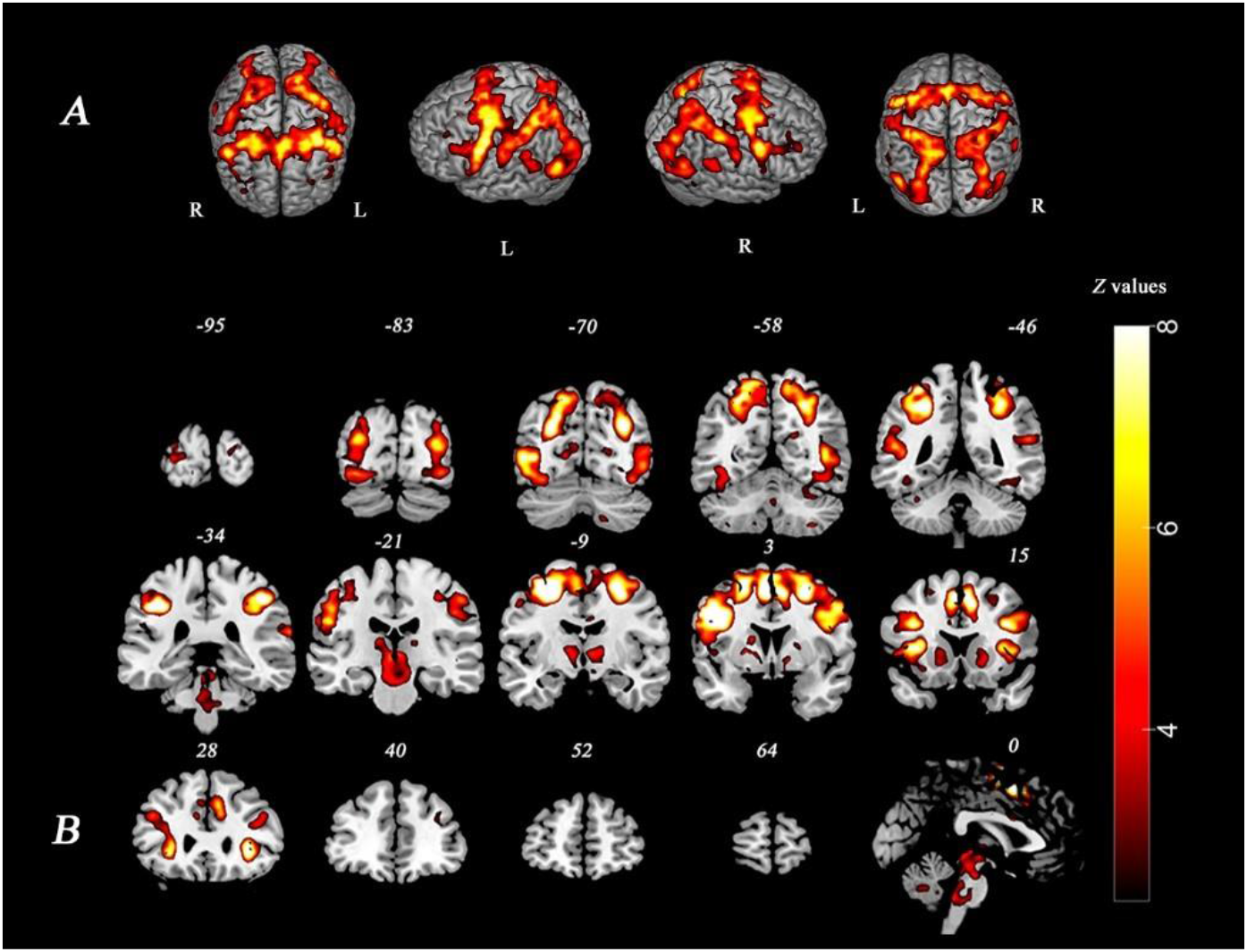
(A) Average fMRI activity across groups and conditions in contrast *all-correct-trials > baseline* at *p*<.05 FWE cluster-level corrected (whole-brain analysis thresholded at Z ≥2.3, rendered view). (B) Average fMRI activity across groups and conditions in contrast *all-correct-trials > baseline* at *p*<.05 FWE cluster-level corrected (whole-brain analysis thresholded at Z ≥ 2.3, slice view).

#### 3.2.2. Region of Interest Analysis

No main effect of Group was obtained for any of the ROIs. The mixed-model ANOVA revealed a main effect of Level of Difficulty (Figure 4) in: (1) left posterior medial frontal cortex, F(3, 84) = 6.93, *p* <.001; (2) right intraparietal sulcus, F(2.13; 59.66) = 5.78, *p =* .004; (3) left inferior frontal gyrus, F(1.95; 54.71) = 3.69, *p =* .03; (4) left intraparietal sulcus, F(2.13; 59.56) = 4.92, *p =* .009; and (5) right caudate, F(3; 84) = 5.12, *p =* .003. However, only left posterior medial frontal cortex and right caudate survived a Bonferroni-correction for 14 comparisons at*p =* .00357. As seen can be seen from Figure 4, fMRI activity in both groups increased in a cubic rather than linear fashion as a function of task difficulty for: (1) left posterior medial frontal cortex, F(1, 28) = 19.71, *p* <.001; (2) right intraparietal sulcus, F(1, 28) = 27.37, *p* <.001; (3) left inferior frontal gyrus,F(1, 28) = 9.95, *p =* .004; and (4) left intraparietal sulcus, F(1, 28) = 17.10, *p*<.001. In addition, pre-HD *=* .004. However, after applying a Bonferroni-correction (*p* = .00357), both groups exhibited a cubic increase of fMRI activity in left posterior medial frontal cortex, as well as in right and left intraparietal sulcus only.

**Figure 4.**
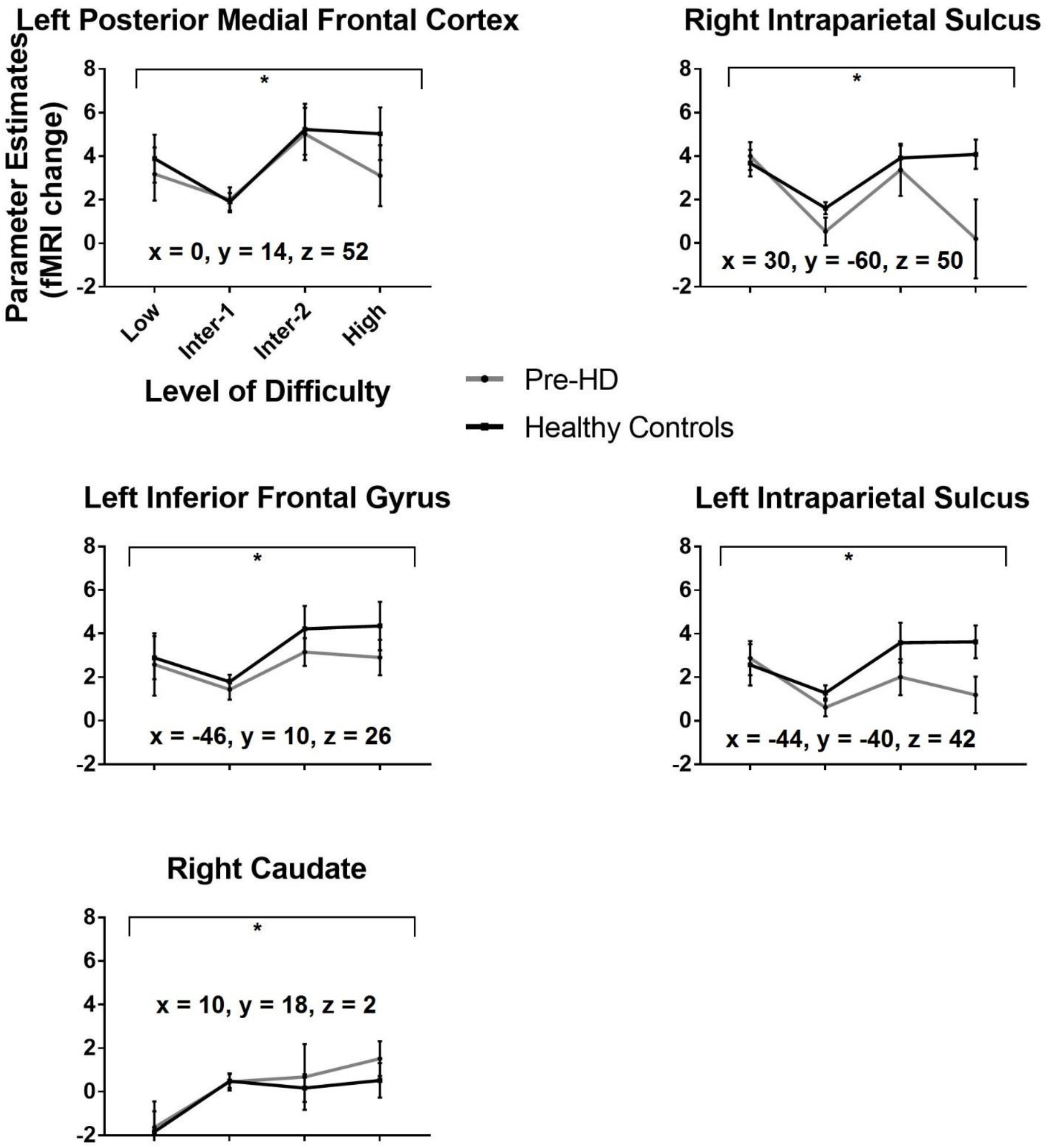
fMRI activity graphs of ROIs showing significant main effect of Level of Difficulty for pre-HD individuals and healthy controls. The grey line corresponds to pre-HD individuals. The black line corresponds to healthy controls. Standard errors are represented in the figure by error bars for each group for each difficulty level. Black asterisk denotes main effect of Level of Difficulty at *p* <.05.

There was no significant Group by Level of Difficulty interaction in any ROI, except in the right intraparietal sulcus, which showed a trend towards significance, F(2.13; 59.66) = 2.78, *p* = .067. This reflects a slight over-recruitment of fMRI activity in pre-HD vs. healthy controls at low level of difficulty and decreased fMRI activity at high level of difficulty.

Consistent with CRUNCH predictions, the *a priori* within-subjects trend analysis revealed a marginally significant linear trend for the Group by Level of Difficulty interaction in left intraparietal sulcus, F(1, 28) = 4.22, *p =* .049, η^2^_p_ =13 (this did not survive a conservative Bonferroni-correction at *p* = .00357) (see Figure 5). Pre-HD individuals exhibited increased fMRI activity at low level of difficulty and decreased fMRI activity at high level of difficulty, compared to healthy controls. Simple effects analyses revealed that pre-HD individuals showed less fMRI activity in left intraparietal sulcus at high level of difficulty, compared to healthy controls, F(1, 28) = 4.63, *p =* .040,η^2^_p_*=* . 14. However, there was no significant difference in fMRI activity between pre-HD and healthy controls in left intraparietal sulcus at low level of difficulty.

**Figure 5.**
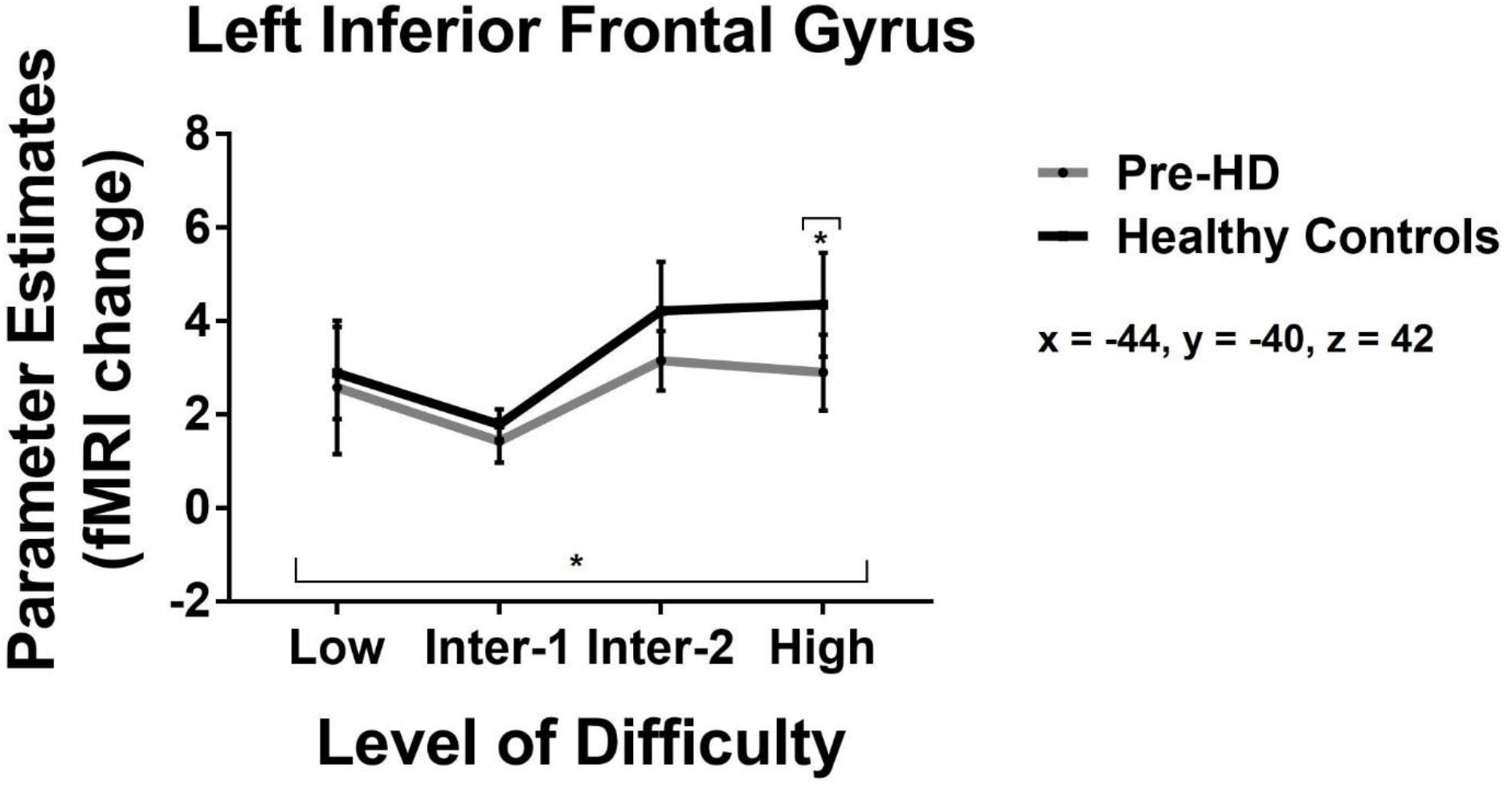
fMRI activity graph of left intraparietal sulcus showing a significant linear trend for the Group by Level of Difficulty interaction for pre-HD individuals and healthy controls. Standard errors are represented in the figure by error bars for each group for each difficulty level. Black asterisks denote a significant linear trend for the Group by Level of Difficulty interaction at *p*<.05 and a significant main effect of Group at high level of difficulty at *p*<.05.

## 4. Discussion

For the first time, we used the CRUNCH model (Reuter-Lorenz & Cappell, 2008), a quantitative model of compensation, to explicitly test compensatory function in pre-HD. We used a novel visuospatial working memory task that parametrically increased level of difficulty (low, intermediate-1, intermediate-2, and high), as is required to test the CRUNCH model predictions (Fabiani, 2012). The behavioural results confirmed that with increasing visuospatial working memory load, RT and error rates increased across both groups. Further, pre-HD individuals performed as effectively as healthy controls at low and intermediate-1 level of task difficulty, and made significantly more errors at intermediate-2 and high level of task difficulty, compared to healthy controls. Consistent with the CRUNCH prediction, fMRI analyses confirmed that pre-HD individuals showed decreased fMRI activity in left intraparietal sulcus at high level of task difficulty, compared to healthy controls. Contrary to the other CRUNCH prediction, we did not observe fMRI over-activation in pre-HD individuals, compared to healthy controls, at lower levels of task difficulty in left intraparietal sulcus, thus providing partial support for validity of CRUNCH in pre-HD. No other ROIs showed the ‘CRUNCH-like’ pattern of fMRI activity in pre-HD, compared to healthy controls.

Visuospatial working memory is one of the first cognitive domains that starts to deteriorate in pre-HD, as evidenced by significantly lower visuospatial working memory capacity than controls (Dumas et al., 2012; Tabrizi et al., 2011). It has been well established that the execution of visuospatial working memory performance relies upon the dorsal striatum (Balleine, Delgado, & Hikosaka, 2007; Lawrence, Watkins, Sahakian, Hodges, & Robbins, 2000; Mizumori, Ragozzino, & Cooper, 2000). Importantly, neurodegenerative anomalies, such as pronounced loss of medium spiny neurons (Albin et al., 1992; Sapp et al., 1997), increased microglial activation (Politis et al., 2011), as well as significant striatal atrophy (Georgiou-Karistianis et al., 2013a) preferentially commence early in the dorsal-striatal network, and are thought to be a critical factor in the development of deficits in visuospatial working memory. In support of this argument, our behavioural data showed that pre-HD individuals exhibited deficits in visuospatial working memory as early as 25 years prior to predicted clinical onset.

While no other HD study has explicitly tested whether CRUNCH could apply in HD; our findings are consistent with Wolf et al. (2007) who also demonstrated a differential effect of task difficulty on the pattern of fMRI activity in pre-HD (*n* = 16) (mean of 19.7 years to disease onset) and healthy controls (*n* =16) during verbal working memory performance. In particular, the pre-HD group showed decreased fMRI activity in left dorsolateral prefrontal cortex at intermediate and high verbal working memory loads, compared with healthy controls (Wolf et al., 2007). In that study, although the pre-HD group performed as effectively as healthy controls on a task at low verbal working memory load, pre-HD did not exhibit increased fMRI activity at low level of difficulty in left dorsolateral prefrontal cortex. Moreover, Kloppel et al. (2009) demonstrated the ‘CRUNCH-like’ pattern of fMRI activity in pre-HD individuals (*n* = 15) (mean of 12.51 years to predicted clinical onset) who showed fMRI over-activation at lower task difficulties in a sequential finger movement task in superior parietal lobule, as well as reduced fMRI activity in this region at higher task difficulties, compared with healthy controls (*n*= 12).

The pre-HD individuals in our study were far from disease onset with a mean of 25.77 years and low disease burden score (mean of 176.33). Previous studies have had samples with higher disease burden scores ≥ 250), with ≤ 15 years to predicted clinical onset (e.g., Enzi et al., 2012; Georgiou-Karistianis et al., 2014; Kloppel et al., 2015; Kloppel et al., 2009). For example, Kloppel et al. (2015) found that only pre-HD individuals close to predicted disease onset (<15 years) showed ‘compensatory’ effects; i.e., increased fMRI activations in right parietal cortex during verbal working memory, compared with healthy controls. Likewise,when performing a response shifting task, pre-HD individuals close to predicted disease onset (mean of 14.8 years) exhibited increased fMRI activity in fronto-temporal regions and basal ganglia, compared with healthy controls (Gray et al., 2013). Based on these findings, it is possible that compensatory fMRI activity occurs during more advanced pre-HD stages when compensatory over-activation may be required as a mechanism to compensate for overt pathology. As such, the pre-HD group in our study may have maintained a ‘healthy-like’ pattern of fMRI activity at lower task demands in the absence of behavioral deficits at low and intermediate-1 level of difficulty because they *may not have required* compensatory over-activation to preserve performance (Nagel et al., 2009).

Furthermore, the pre-HD group in our study exhibited significantly reduced fMRI activity in left intraparietal sulcus at high level of difficulty compared to heathy controls, who instead showed increased fMRI activity, as predicted by CRUNCH. Our findings are consistent with a number of studies which previously showed that fMRI activity decreased with higher task demands in pre-HD individuals during verbal working memory (Wolf et al., 2007), spatial working memory (Georgiou-Karistianis et al., 2014), and motor performance (Kloppel et al. 2009).

Intraparietal sulcus plays a critical role in maintenance and manipulation of information in visuospatial working memory (Bray, Almas, Arnold, Iaria, & MacQueen, 2015; Linden et al., 2003; Papadopoulos, Sforazzini, Egan, & Jamadar, 2017), and it has been shown that fMRI activity in this region increases in response to memory load during visuospatial working memory (Harrison, Jolicoeur, & Marois, 2010). As such, it is possible that pre-HD individuals showed a decline in fMRI activity in left intraparietal sulcus at high level of task difficulty, as well as a performance drop at intermediate-2 and high level of difficulty on the behavioral level, because they reached their visuospatial working memory capacity (the ‘CRUNCH’ point), compared to healthy controls. Our findings suggest that manipulation of task difficulty can differentially affect the pattern of fMRI activity in left intraparietal sulcus in pre-HD vs. healthy controls, supporting the CRUNCH hypothesis.

Our study had some limitations. We acknowledge that while our sample size is consistent with the standard in the pre-HD literature (e.g., Enzi et al., 2012; Kloppel et al., 2009; Malejko et al., 2014; Wolf et al., 2007), it is small (*n* = 15, pre-HD; *n* = 15, healthy controls). Additionally, pre-HD individuals were very far from disease onset and had no overt pathology, and perhaps this limited the need for compensatory activity. Lastly, fMRI activity at the lowest memory load was higher than activity at intermediate-1 memory load in right and left intraparietal sulcus, left posterior medial frontal cortex and left inferior frontal gyrus. This conflicts with the behavioural results, which showed lower RT and error rate with increased task difficulty, as expected. This result may be analogous to a neural ‘restart cost’, where the first few trials of a new task incur a cost relative to later trials (Monsell, 2003). The crucial comparison here, however is the differential effect between groups; as such this finding does not affect the ability to interpret the results.

To conclude, our study has demonstrated partial support for the validity of CRUNCH model in pre-HD. In light of our results we suggest that CRUNCH, developed in the context of the normal healthy ageing, may not be the most sensitive and robust model to detect compensatory mechanisms early in a pathological disease such as HD. Further research with larger and more heterogeneous (e.g., structural pathology, disease burden score) pre-HD samples is required to clarify whether CRUNCH can be used to competently characterise compensation in pre-HD.

## Ethical statement

The study complies with the national ethical research guidelines in Australia. The study was approved by the Monash University and Melbourne Health Human Research Ethics Committees.

## Informed consent

Written informed consent was obtained from each participant in accordance with the Helsinki Declaration.

## Acknowledgement

Jamadar is supported by an Australian Research Council (ARC) Discovery Early Career Research Award (DE150100406) and the ARC Centre of Excellence for Integrative Brain Function (CE140100007).Poudel is supported by the Hereditary Disease Foundation USA Fellowship and Huntington’s Disease Association (NSW)

